## Supporting Information for "Multiscale modeling of presynaptic dynamics from molecular to mesoscale"

### Estimating $[Ca^{2+}]_i$ from Collision Events

Because of the quantized nature and low concentration of $Ca^{2+}$ ions in the presynaptic space, calculating the instantaneous local calcium concentration just around the SNARE complex of a single docked vesicle is nontrivial in MCell. Instead, we use effector tiles, small virtual surfaces in the presynaptic space of the MCell environment, to estimate local concentration from the frequency of calcium ions passing through them. This section provides a derivation of average $[Ca^{2+}]_i$ from the number of "hits", $N_H$, of calcium ions through the effector tile surface.

For a particle diffusing by Brownian motion in $d$ dimensions, the probability density function $\rho$ of the particle's displacement $r$ from its initial position after a time $\Delta t$ is equal to

$$\rho(r, \Delta t) = \frac{1}{\pi^{d/2}\lambda^d} \mathrm{e}^{-r^2/\lambda^2}, \tag{28}$$

where $\lambda$ is a diffusion length parameter that depends on the diffusion constant and time step. Since we are dealing with calcium, we use

$$\lambda_{Ca} = \sqrt{4D_{Ca}\Delta t}, \tag{29}$$

where $D_{Ca} = 220\ \mu\mathrm{m}^2/\mathrm{sec}$ is the calcium diffusion constant [35]. More directly useful, though, is the average step length along any given axis, in particular, along the component perpendicular to the calcium-detecting surface:

$$\bar{l}_{\perp} = \frac{\lambda_{Ca}}{\sqrt{\pi}} = \sqrt{\frac{4D_{Ca}\Delta t}{\pi}}. \tag{30}$$

Thinking about the effective volume near the effector tile, the expected number of hits of particles through the surface from either side during the interval $\Delta t$ becomes

$$N_H = N_A \bar{l}_\perp A_{ET} [\mathrm{Ca}^{2+}]_i, \tag{31}$$

where $N_A$ is Avogadro's number and $A_{ET}$ is the area of the effector tile. Solving for concentration,

$$[\mathrm{Ca}^{2+}]_i = \frac{N_H}{N_A \bar{l}_\perp A_{ET}}. \tag{32}$$

Now, the average concentration from the start of the simulation until time $t$ becomes

$$c(t) = \frac{N_H(t)}{N_A \bar{l}_\perp A_{ET}} \cdot \frac{\Delta t}{t}, \tag{33}$$

where $N_H(t)$ is the running total number of hits. To find the average $\mathrm{Ca}^{2+}$ concentration over an arbitrary interval $[t_i, t_j]$:

$$\langle [\mathrm{Ca}^{2+}]_i([t_i, t_j]) \rangle = \frac{t_j c(t_j) - t_i c(t_i)}{t_j - t_i}. \tag{34}$$

For each spike train used as input to the simulation, we averaged the instantaneous local active zone calcium concentration over 2000 trials in time steps of 0.1 ms.

### Chemical Kinetics of Calcium Channels, Buffers, and Pumps

The kinetic schemes and kinetic rate constants for the voltage-dependent calcium channel (VDCC), calbindin (CB), and plasma membrane $\mathrm{Ca}^{2+}$-ATPase (PMCA) pump models used for this paper, along with their associated references, are shown below in S1 Fig and S1 Table.

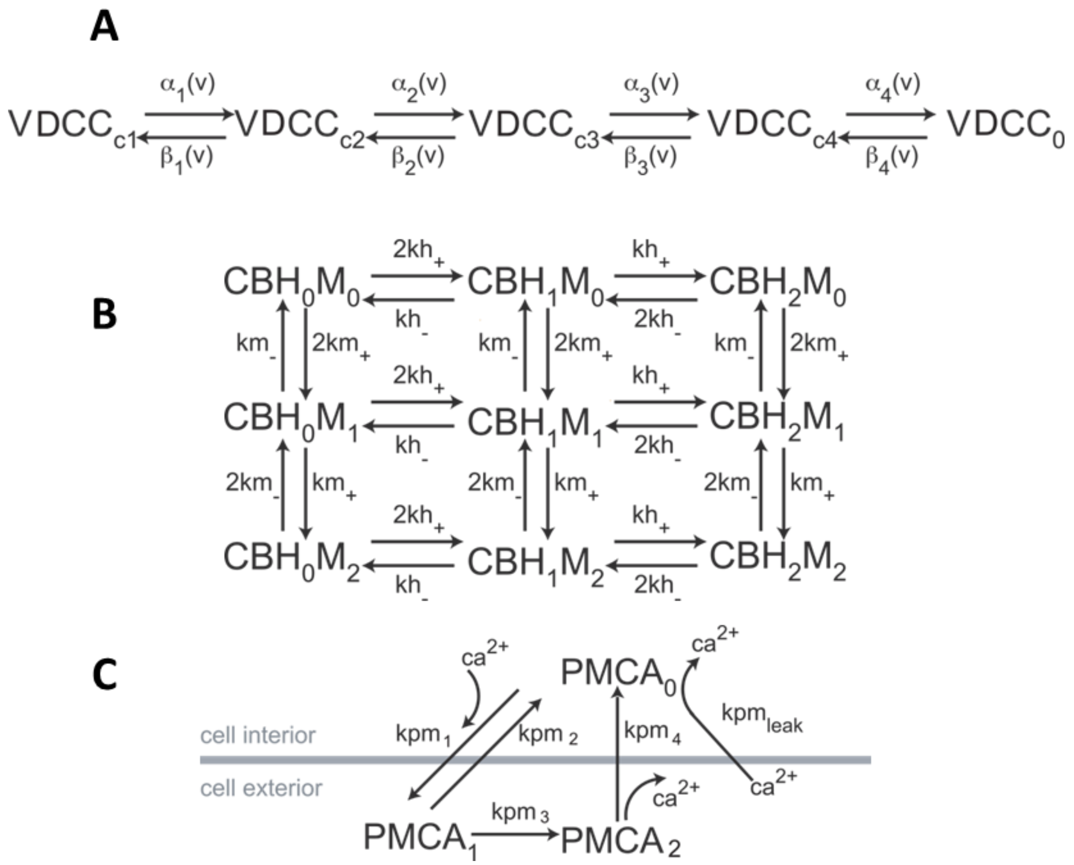


**S1 Fig. State Diagrams for VDCC, Calbindin, and PMCA.**

All diagrams reproduced with permission from Nadkarni et al. [35]. A: VDCC state transition model adapted from Bischofberger et al.[68]. Transition rates $\alpha_{ij}$ and $\beta_{ji}$ depend on membrane potential v. B: State transitions for calbindin (CB) at high-affinity (H) and medium-affinity (M) $Ca^{2+}$-binding sites. On rates ($kh_+$ and $km_+$) are proportional to $[Ca^{2+}]_i$. C: PMCA pump state diagram with $Ca^{2+}$ interactions depicted on the relative side of the membrane. $Ca^{2+}$ leakage occurs only in state $PMCA_0$. Association rate $kpm_1$ is proportional to $[Ca^{2+}]_i$.

**S1 Table. Parameter Values for VDCC, Calbindin, and PMCA.**

Table adapted from [35]. VDCC rates follow $\alpha_i(v)=\alpha_{i0}\exp(v/v_i)$ and $\beta_i(v)=\beta_{i0}\exp(-v/v_i)$. VDCC parameters values adapted from [68]. Calbindin parameter values adapted from [77]. PMCA parameter values adapted from [78].

| Parameter | Value |
|---|---|
| **VDCC - [68]** | |
| $\alpha_{10}, \alpha_{20}, \alpha_{30}, \alpha_{40}$ | 4.04, 6.70, 4.39, 17.33 $\mathrm{ms}^{-1}$ |
| $\beta_{10}, \beta_{20}, \beta_{30}, \beta_{40}$ | 2.88, 6.30, 8.16, 1.84 $\mathrm{ms}^{-1}$ |
| $v_1, v_2, v_3, v_4$ | 49.14, 42.08, 55.31, 26.55 $\mathrm{mV}$ |
| **Calbindin-D28k - [77]** | |
| $kh_+$ | $5.5 \times 10^6\ \mathrm{M}^{-1}\mathrm{s}^{-1}$ |
| $kh_-$ | $2.6\ \mathrm{s}^{-1}$ |
| $km_+$ | $4.35 \times 10^7\ \mathrm{M}^{-1}\mathrm{s}^{-1}$ |
| $km_-$ | $35.8\ \mathrm{s}^{-1}$ |
| **PMCA - [35, 78]** | |
| $kpm_1$ | $1.5 \times 10^8\ \mathrm{M}^{-1}\mathrm{s}^{-1}$ |
| $kpm_2$ | $20\ \mathrm{s}^{-1}$ |
| $kpm_3$ | $100\ \mathrm{s}^{-1}$ |
| $kpm_4$ | $1.0 \times 10^5\ \mathrm{s}^{-1}$ |
| $kpm_{leak}$ | $12.264\ \mathrm{s}^{-1}$ |

In response to an action potential stimulus, voltage-dependent $Ca^{2+}$ channels (VDCCs) transition stochastically to an open state, through which $Ca^{2+}$ ions may enter the axon down a sharp electrochemical gradient [68, 126]. Because this process does not depend on diffusion, a deterministic simulation of state probabilities can perfectly capture the shape of the histogram of $Ca^{2+}$ influx rate averaged over infinite trials, as in S2 Fig. Notice that the rate of influx rises to a peak and returns completely to baseline within a span of about 2 ms, so any spike-evoked vesicle fusion after this initial influx is due entirely to internal dynamics as $Ca^{2+}$ diffuses, interacts with the buffer and $Ca^{2+}$ sensors, and vacates through the pumps.

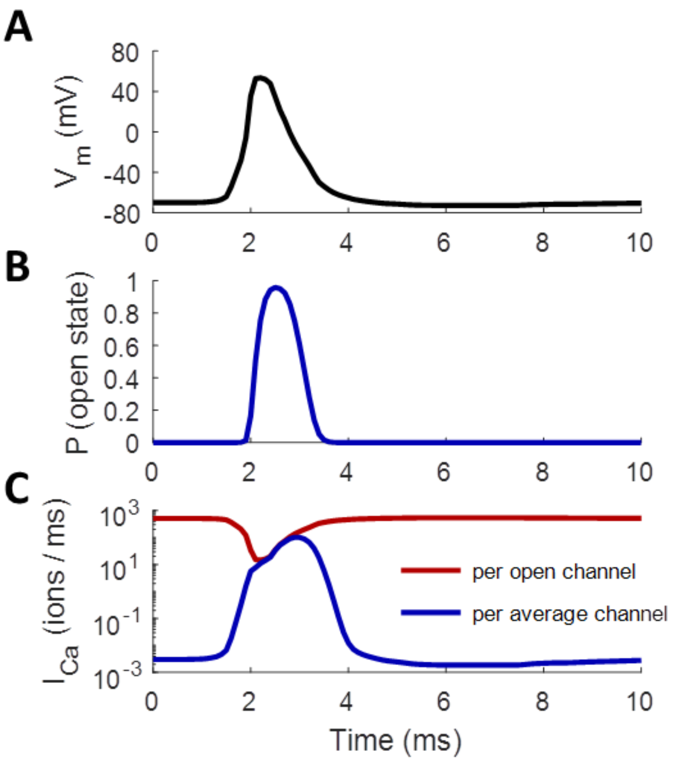


**S2 Fig. Action-Potential-Evoked $Ca^{2+}$ Current.**

A: Action-potential-like waveform applied to axon. B: Probability of a single VDCC being in the open state in response to the action potential in panel A increases from about $10^{-5}$ to around 96% during the spike before quickly shutting off; computed from deterministic simulation of state probabilities. C: Rate of $Ca^{2+}$ influx through a single, pathologically open channel (red) and through a typical channel (blue), whose probability of being open follows B.

Of course, the existence of a nonzero $[Ca^{2+}]_{i0}$ implies that the $Ca^{2+}$-sensors of the SNARE complex will induce vesicle fusion at some finite, if extremely slow, rate. At very low concentrations, this would require anywhere from many thousands to many trillions of trials to build up sufficiently informative release histograms. Instead, we reran the deterministic model at constant values of $[Ca^{2+}]_i$ with no $Ca^{2+}$ spike and measured the steady-state release rates after 10 seconds of simulated time (S3 Fig). Perhaps unsurprisingly, the spontaneous release rates grow in proportion to the 5$^{th}$ (2$^{nd}$) power of $[Ca^{2+}]_{i0}$ for synchronous (asynchronous) release, according to the number of $Ca^{2+}$ ions needed to bind before the synaptotagmin can initiate fusion. At very high $[Ca^{2+}]_{i0}$, though, the release rates saturate to $\gamma_S$ and $\gamma_A$ (see Table 1) as the probability of being in the releasable state approaches one.

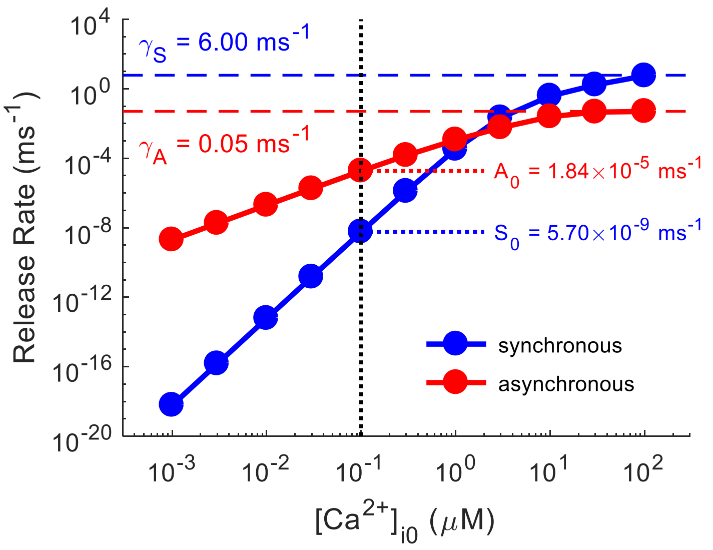


**S3 Fig. Spontaneous Rates of Vesicle Fusion Increase with $[Ca^{2+}]_{i0}$.**

For small $[Ca^{2+}]_{i0}$, $S_0=k_S\cdot([Ca^{2+}]_{i0})^5$ and $A_0=k_A\cdot([Ca^{2+}]_{i0})^2$, where $k_S\approx6\times10^{-4}$ ms$^{-1}\cdot\mu M^{-5}$ and $k_A\approx2\times10^{-3}$ ms$^{-1}\cdot\mu M^{-2}$. As $[Ca^{2+}]_{i0}\longrightarrow\infty$, $S_0\longrightarrow\gamma_S$ and $A_0\longrightarrow\gamma_A$. Values for $S_0$ and $A_0$ at $[Ca^{2+}]_{i0}$=100 nM, which is used throughout most of this paper, are pointed out for reference.

### Effects of Buffer and Spatial Modeling on Release Dynamics

Running these simulations in MCell, rather than as a much simpler well-mixed model, was essential for capturing both distance-dependent effects and temporal features of the $Ca^{2+}$ waveform. The well-mixed assumption, which ignores diffusion and treats all chemical processes as occurring at the same point in space, does not hold at the spatial and temporal scales of interest in the synapse [46, 47]. As seen in Fig 2C, peak $Ca^{2+}$ drops precipitously even over fractions of a micron away from the VDCC cluster, and the shape of the response changes dramatically over this same scale, transitioning from a predominantly synchronous to a predominantly asynchronous profile. These trends, elucidated by the spatial MCell simulation, are completely absent in the space-less well-mixed simulation (maroon curves, S4 Fig), even when all other aspects of the model remain the same, such as the number of VDCCs, calbindin buffer molecules, and PMCA pumps and the set of all state transitions for each molecular species. Note also from S4 Fig A that the transition in time from the fast synchronous component to the extended asynchronous component is much sharper in the case without space. The extra $Ca^{2+}$ decay component arises from local saturation effects. After the initial rapid influx, the calbindin buffer immediately around the VDCC cluster becomes saturated, causing the high free $Ca^{2+}$ that remains to overwhelm the PMCA pumps' ability to evacuate it from the area. The pumps remove it at a constant maximum rate, leading to a short linear decay only evident very near the VDCCs (yellow traces, S4 Fig A) or when all calbindin is removed from the simulation (S4 Fig B). Such local saturation effects do not appear in the well-mixed case because all buffer molecules and pumps are simultaneously available to all the free $Ca^{2+}$ ions. Thus, in light of all these effects, the spatial MCell model is crucial for the task of properly characterizing the $Ca^{2+}$ transient in the synapse.

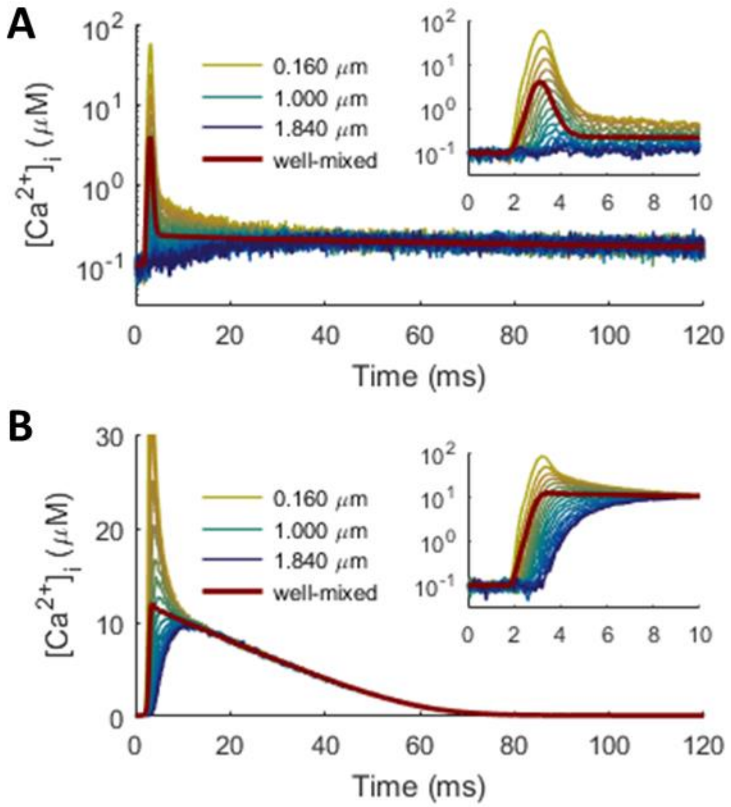


**S4 Fig. Spatial Modeling Important for Capturing Fine-Grain Features of $Ca^{2+}$ Transients.** Color scheme identical to that used in Fig 2: yellow to blue represent proximal to distal $Ca^{2+}$ sensors. A: $[Ca^{2+}]_i$ measured at increasing distance from VDCC source (yellow to blue), with well-mixed approximation overlaid for comparison (maroon). Inset focuses on shorter time scale. B: Profiles with calbindin removed from MCell (yellow to blue) and well-mixed model (maroon). Note that peak $[Ca^{2+}]_i$ for the most proximal case extends up to 81 μM, but is cut off for clarity.

Most neurotransmitter release occurs within a sharp window after an action potential stimulus [127-130]. The presence of the $Ca^{2+}$ buffer calbindin plays an instrumental role in this by rapidly removing most of the free $Ca^{2+}$ and then slowly releasing it over an extended period at a rate that the active PMCA pumps can handle. This action significantly tightens the window for Syt-1-mediated synchronous release [40, 131] while also extending the time window for Syt-7-mediated asynchronous release. Without a buffer, however, the free $[Ca^{2+}]_i$ does not drop off immediately but decays linearly toward baseline over a few tens of milliseconds, saturating the capacity of the PMCA pumps to remove the ions (S4 Fig B, S5 Fig A,C). Thus, removing calbindin from the simulations both amplifies synchronous release in a time window near the spike and suppresses asynchronous release long after the stimulus (S5 Fig B,D). This agrees with experimental evidence that endogenous $Ca^{2+}$ buffers limit the rate of synchronous synaptic release [131].

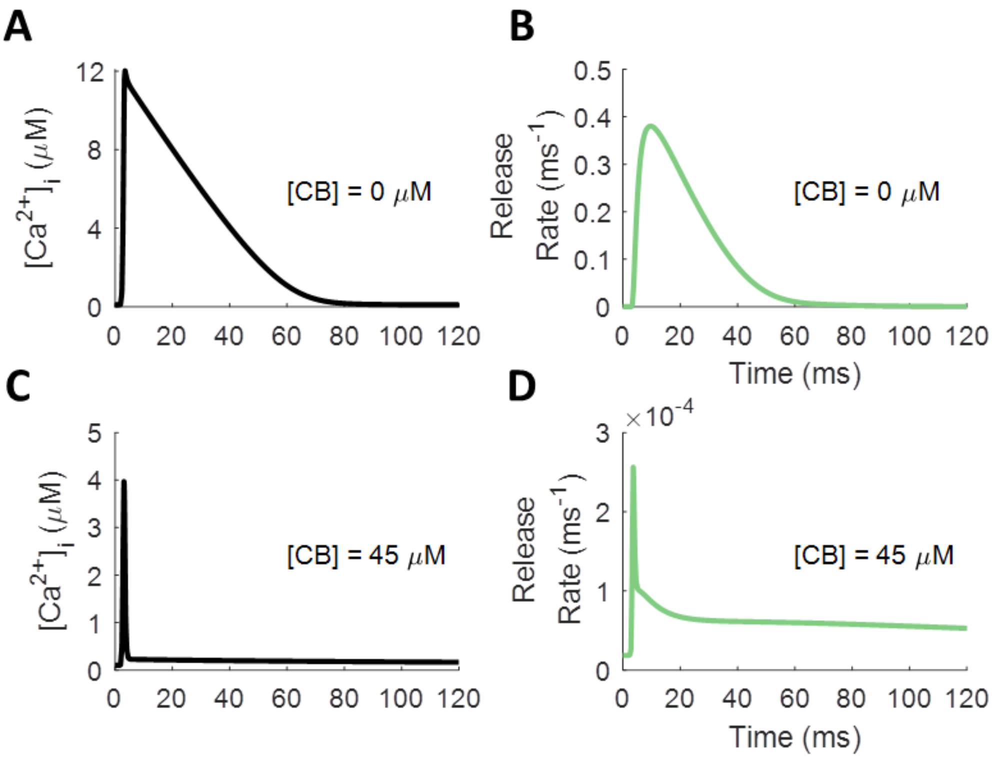


**S5 Fig. Effect of Calbindin Buffer on Spike-Evoked $Ca^{2+}$ Profile and Release Rates.**
Action-potential-like stimulus delivered to model axon starting at 0 ms. Diffusion is assumed to be instantaneous, and molecular state probabilities are tracked deterministically over time. A: Free $[Ca^{2+}]_i$ with no calbindin buffer decays linearly with time due to saturation of PMCA pumps. B: Syt-1/7-mediated release rates are large but short-lived in response to unbuffered $Ca^{2+}$. C: Free $[Ca^{2+}]_i$ with calbindin added to the axon has much smaller magnitude and much narrower peak but has much longer tail. D: Vesicle release in response to buffered $Ca^{2+}$ is much less pronounced. The calbindin buffer reduces the rate of synchronous transmission but extends the window for pronounced asynchronous transmission.

After obtaining the distance-dependent $Ca^{2+}$ traces, we could use them to see how the rate of release changes with distance. Using the above-measured $Ca^{2+}$ traces as input to the deterministic Markov model of Syt-1/7, we once again calculated the instantaneous rates of spike-evoked release for single vesicles at increasing distances. As expected, the single-vesicle probability of release decays with distance until it reaches a distance-independent baseline level (S6 Fig), although this occurs differently for the synchronous and asynchronous mechanisms.

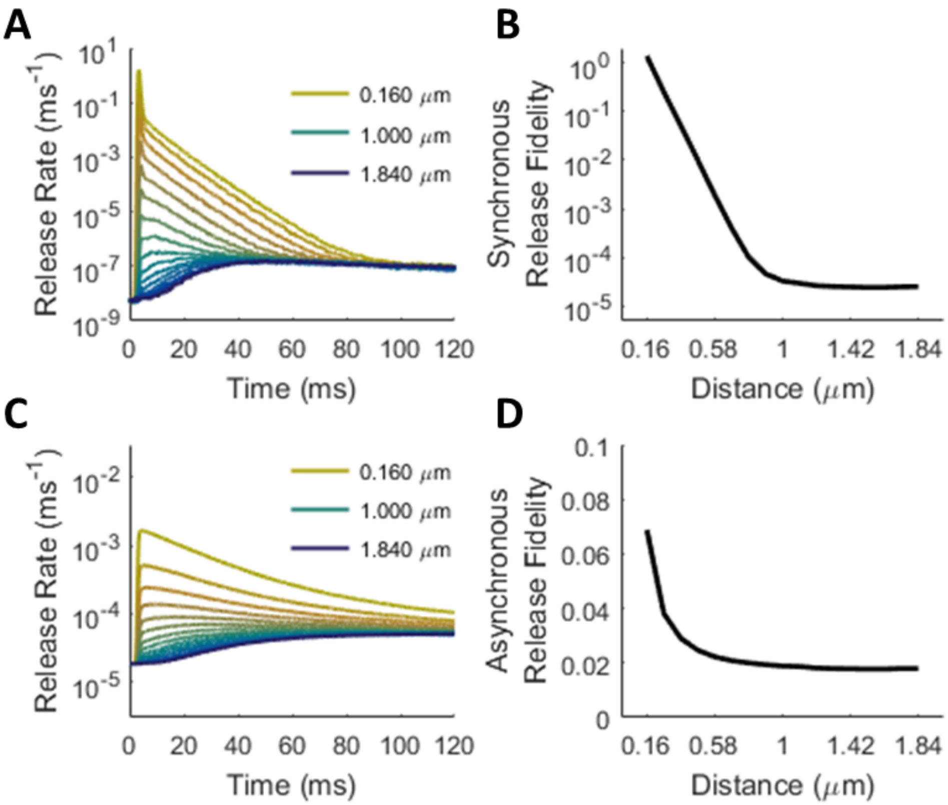


**S6 Fig. Synchronous and Asynchronous Release Rates Decrease with Distance from the $Ca^{2+}$ Source.**

Color scheme identical to that used in Fig 2 and S4 Fig: yellow to blue represent proximal to distal $Ca^{2+}$ sensors. A: Synchronous release rate. B: Integrated probability of synchronous release falls off nearly exponentially with distance to a baseline level. C: Asynchronous release rates. D: Integrated probability of asynchronous release also decays with distance to some baseline, but not exponentially.

To account for the change in release profiles mathematically, we ran a fitting algorithm on each profile, exploring the space of values both for the magnitude of each component of release ($P_c$ in Eq (1) and (3)) and for the temporal filter parameters ($k_c$, $\mu_c$, and $\sigma_c$ in Eq (3) and (4)). We assumed that the time constants of release rate decay ($\tau_c$) remained the same for the release histograms at all distances and that any changes in the size or shape in the histograms are due to depleted levels of $[Ca^{2+}]_i$ and to increasing delays for $Ca^{2+}$ ions to reach the sensors. Accordingly, we expected to see the $P_c$ values decay with distance as $Ca^{2+}$ is dissipated, sequestered, and removed; the $k_c$ values to slow down as the limiting delay grows with distance; and the values of $\mu_c$ and $\sigma_c$ to increase somewhat due to greater numbers of potential interactions before the $Ca^{2+}$ ions complete their traversal. The fitting algorithm produced sets of parameters at each location in the synapse that generally followed these trends (S7 Fig C,D), although the noise in the data and the very high dimensionality of the problem prevented smooth trends from being ascertained.

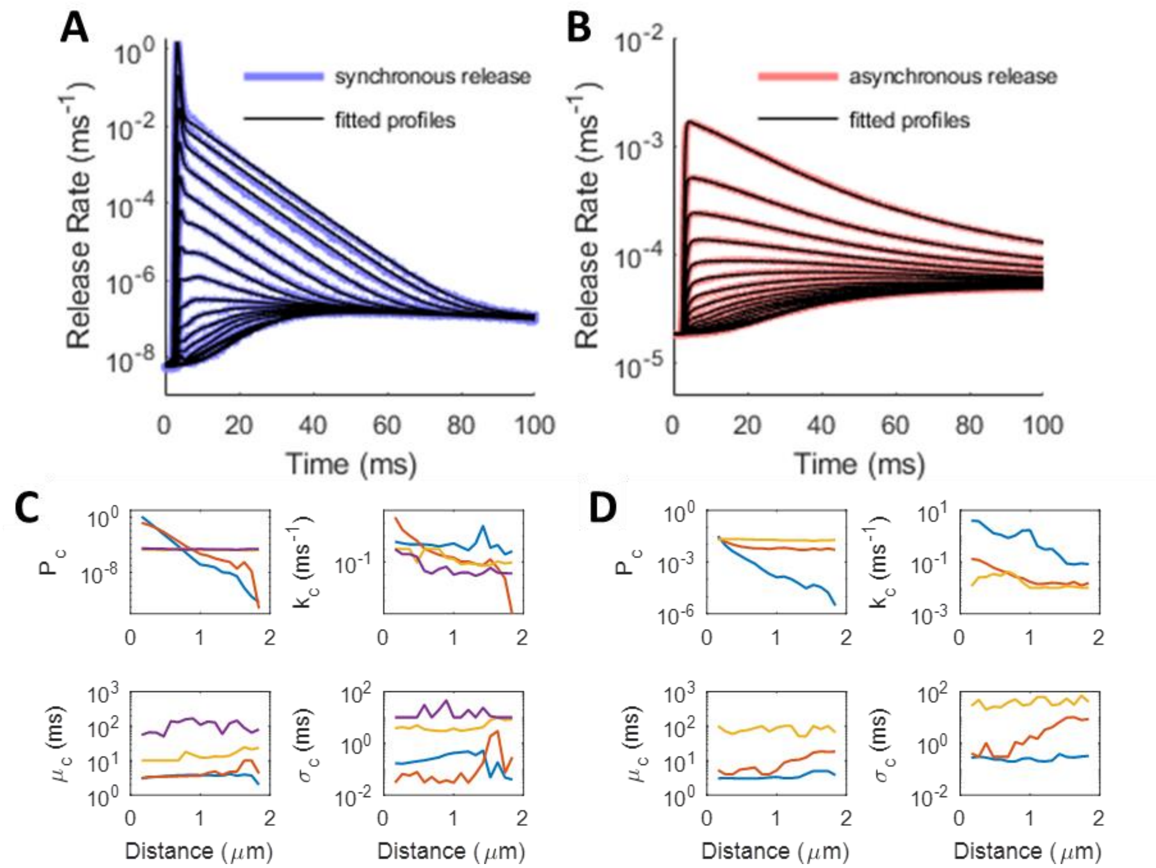


**S7 Fig. Parametric Fits to Release Histogram Profiles at Increasing Distance from the $Ca^{2+}$ Source.**

A, B: Fitted release profiles (black) imposed over the true histograms for synchronous (A, blue) and asynchronous (B, red). C: Parameter values as a function of distance for synchronous release. D: The same for asynchronous release.

### Applying Release Start Time Filter to Release Rate Profiles

The release-start-time filter introduced in Eq (3) and (4) follows an ex-Gaussian distribution, $a(t; k_c, \mu_c, \sigma_c)$, representing trial-to-trial variation in the start time for spike-evoked release due to the stochasticity of buffered diffusion. This can be treated as adding an exponentially distributed random delay with rate parameter $k$ and a normally distributed random delay with mean $\mu$ and standard deviation $\sigma$ to the spike time at $t = 0$, removing subscripts for simplicity. Applying this filter to a release profile component $r(t) = P/\tau \left(\mathrm{e}^{-t/\tau}\mathrm{u}(t)\right)$ from Eq (1) and (3) requires performing a convolution operation as shown below:

$$\begin{aligned} r(t) &= \frac{P}{\tau}\left(\mathrm{e}^{-t/\tau}\mathrm{u}(t)\right) * a(t; k, \mu, \sigma) \\ &= \frac{P}{\tau}\left(\mathrm{e}^{-t/\tau}\mathrm{u}(t)\right) * \left(k\mathrm{e}^{-kt}\mathrm{u}(t)\right) * \left(\frac{1}{\sigma\sqrt{2\pi}} e^{-\frac{(t-\mu)^2}{2\sigma^2}}\right) \\ &= \left(P\frac{k}{k\tau - 1}\left(\mathrm{e}^{-t/\tau} - \mathrm{e}^{-kt}\right)\mathrm{u}(t)\right) * \left(\frac{1}{\sigma\sqrt{2\pi}} e^{-\frac{(t-\mu)^2}{2\sigma^2}}\right). \end{aligned} \tag{35}$$

Stopping here and replacing the Gaussian component with a delta function by letting $\sigma \to 0$ yields

$$r(t) = P\frac{k}{k\tau - 1}\left(\mathrm{e}^{-(t-\mu)/\tau} - \mathrm{e}^{-k(t-\mu)}\right)\mathrm{u}(t - \mu), \tag{36}$$

which includes both an initial phase where release rate ramps up after $t = \mu$ and a decay phase where release rate falls off exponentially. Note that the area under the curve, and thus the probability of release, remains the same. For $\sigma > 0$, the final form of the release component looks like

$$r(t) = P\frac{k}{k\tau - 1}\left(\mathrm{e}^{-\left(t-\left(\mu+\frac{\sigma^2}{2\tau}\right)\right)/\tau}\Phi\left(\frac{t-(\mu+\sigma^2/\tau)}{\sigma}\right) - \mathrm{e}^{-k\left(t-\left(\mu+\frac{\sigma^2}{2}k\right)\right)}\Phi\left(\frac{t-(\mu+\sigma^2 k)}{\sigma}\right)\right), \tag{37}$$

which basically just adds a little extra rightward temporal shift and smooths out the corner in the profile shape, due to replacing the step function of Eq (36) with the CDFs of two normal distributions. Fig 8 A-C shows how this filter affects the shape of a release profile component.

### Facilitation Nonlinearities

Release probability increases from the start of an action potential to its peak and from one spike to the next because of the accumulation of $Ca^{2+}$ on the sensor in the SNARE complex. Even when not enough $Ca^{2+}$ has accumulated to trigger vesicle fusion on the first spike, it can still increase the probability of reaching the releasable state after subsequent spikes. As can be seen in S8 Fig, $Ca^{2+}$ entry from one spike can predispose the distribution of bound states of the sensor to trigger release with greater alacrity on subsequent spikes.

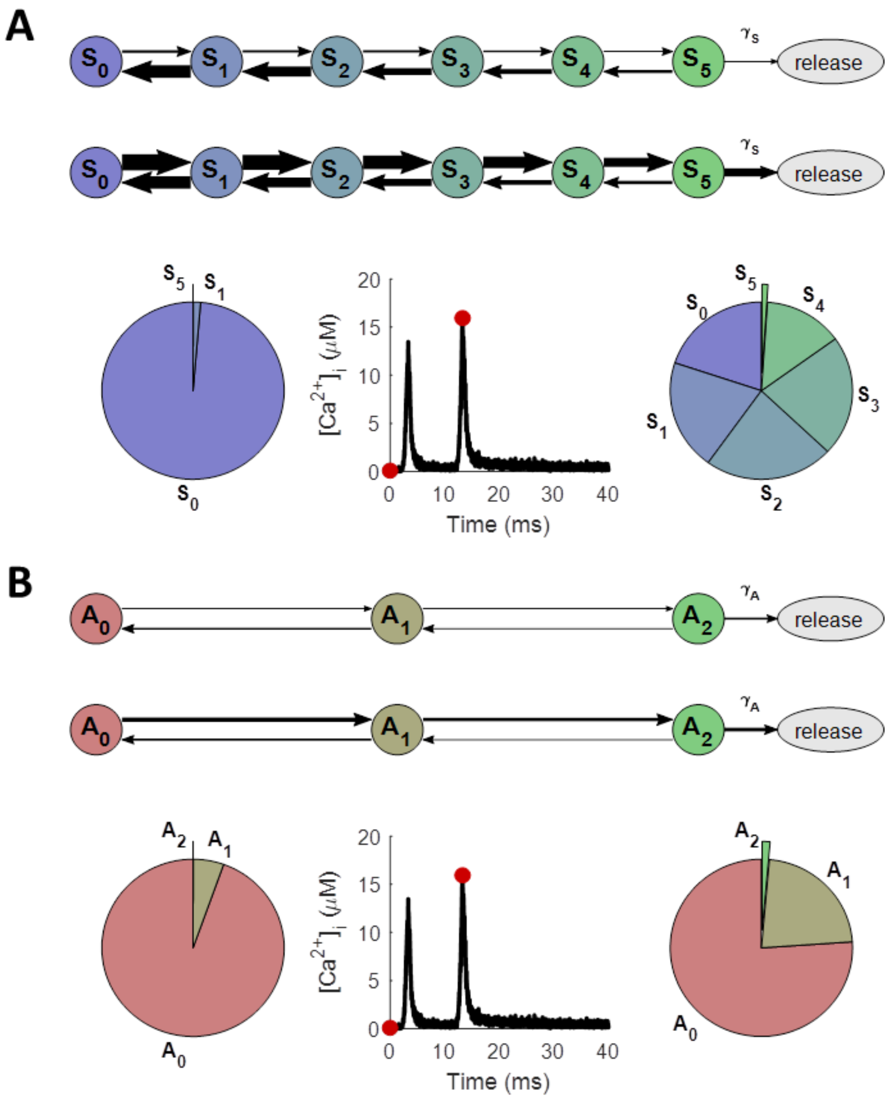


**S8 Fig. Change in the Balance of Binding Kinetics and Internal State Distribution of $Ca^{2+}$ Sensor with Spike History.**

State diagrams the same as shown in Fig 3. A: Synchronous state diagrams. At baseline $[Ca^{2+}]_i$ (first red dot), unbinding kinetics (left arrows) overpower binding (right arrows), biasing Syt-1 toward unbound state ($S_0$; top diagram), with almost no probability of having any $Ca^{2+}$ ions bound before an action potential (left pie chart). During peak $Ca^{2+}$ influx (second red dot), binding rates (thicker right arrows) overpower unbinding, biasing Syt-1 toward its fully-bound releasable state ($S_5$; lower diagram), with much greater probability of having at least some $Ca^{2+}$ bound (right pie chart). B: The same for asynchronous release with Syt-7, whose releasable state requires two $Ca^{2+}$ ions bound ($A_2$). Slower kinetics lead to only slight bias in favor of binding during an action potential (slightly thicker right arrows in lower diagram), leading to miniscule increase in probability of being in the releasable state on later spikes (right pie chart). Release becomes more probable on subsequent spikes because previous activity has pushed synaptotagmin into higher-bound states, making reaching the releasable state easier.

Simulations with the MCell model demonstrate how nonlinear binding cooperativity in the $Ca^{2+}$ sensors induces facilitation in excess of what would be expected from cytoplasmic $Ca^{2+}$ buildup alone. S9 Fig shows how the combined release rate from synchronous and asynchronous release mechanisms (dark green: spike ramp; light green: probe spikes of different trains) grows far more quickly than does spike-evoked $[Ca^{2+}]_i$ (black/gray). Thus, the magnitude of facilitation may be nonlinear due to the internal binding kinetics of the synaptotagmin.

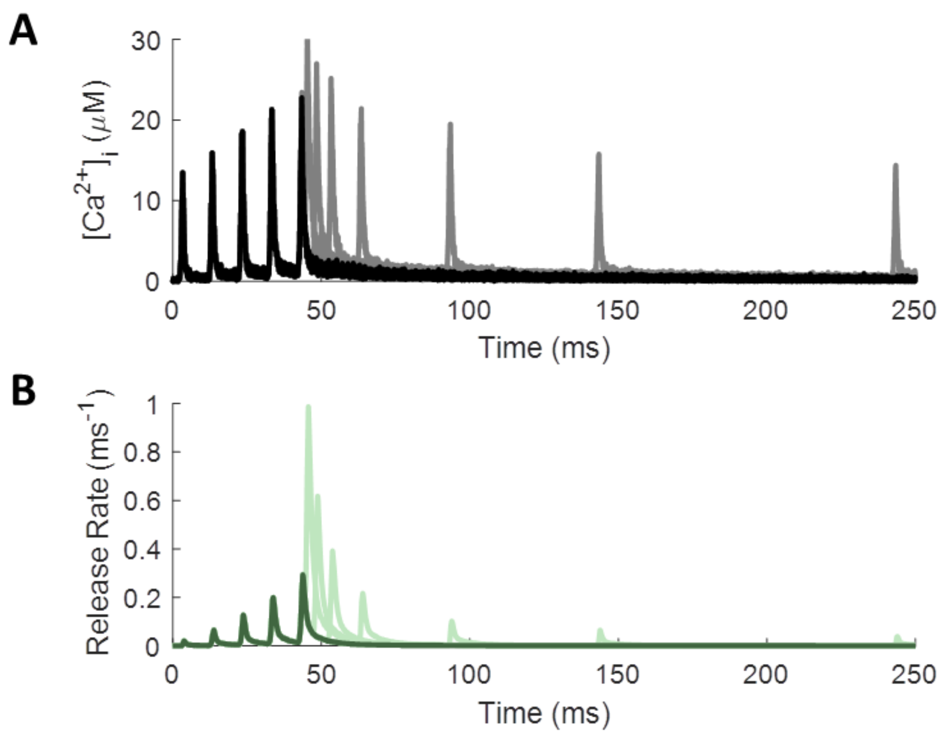


**S9 Fig. Empirical Facilitation in Release Probability is a Nonlinear Function of Spike History and $Ca^{2+}$ Buildup.**

A: $[Ca^{2+}]_i$ and release rate in response to a 5-spike ramp stimulus with a 10-ms ISI (black and dark green), followed by a single probe spike at increasing delay from the end of the ramp (gray and light green; multiple cases overlaid on the same plot). Release rate grows much faster than $Ca^{2+}$ buildup can account for.

As described in Methods, we explored facilitation for 136 unique spike trains, each composed of a constant-frequency spike ramp followed by a single probe spike at increasing interspike intervals (ISI). As an example, S9 Fig overlays multiple spike trains, each with a spike ramp of 5 spikes with 10-ms ISI (dark colors) and each with a separate probe spike at exponentially increasing ISI (light colors). To gain an intuition of how facilitation varies across different spike histories, we calculated the integrated release magnitude of the final spike of each train, according to

$$P(n) = \int_{t_{sn}}^{\infty} (r^*(t - t_{sn}) - r_0)dt = \sum_{c=1}^{N} P_c(n), \tag{38}$$

where $t_{sn}$ is the time of spike n, $r^*(\cdot)$ is the empirical release rate function after this spike, given that no further activity occurs, and $r_0$ is the spontaneous release rate. The empirical facilitation factor is simply the ratio of integrated release magnitude on spike $n$ to that on spike 0:

$$F(n) = \frac{P(n)}{P(0)}, \tag{39}$$

Note that this empirical facilitation factor applies to the sum of all release components and does not correspond to any one component specifically. As can be seen in S10 Fig, empirical facilitation increases along a constant-frequency spike ramp and diminishes thereafter for increasing ISI of the following probe spike back toward baseline. However, the level of facilitation is not a simple function of the most recent activity but depends on the rate of stimulation prior to the last spike.

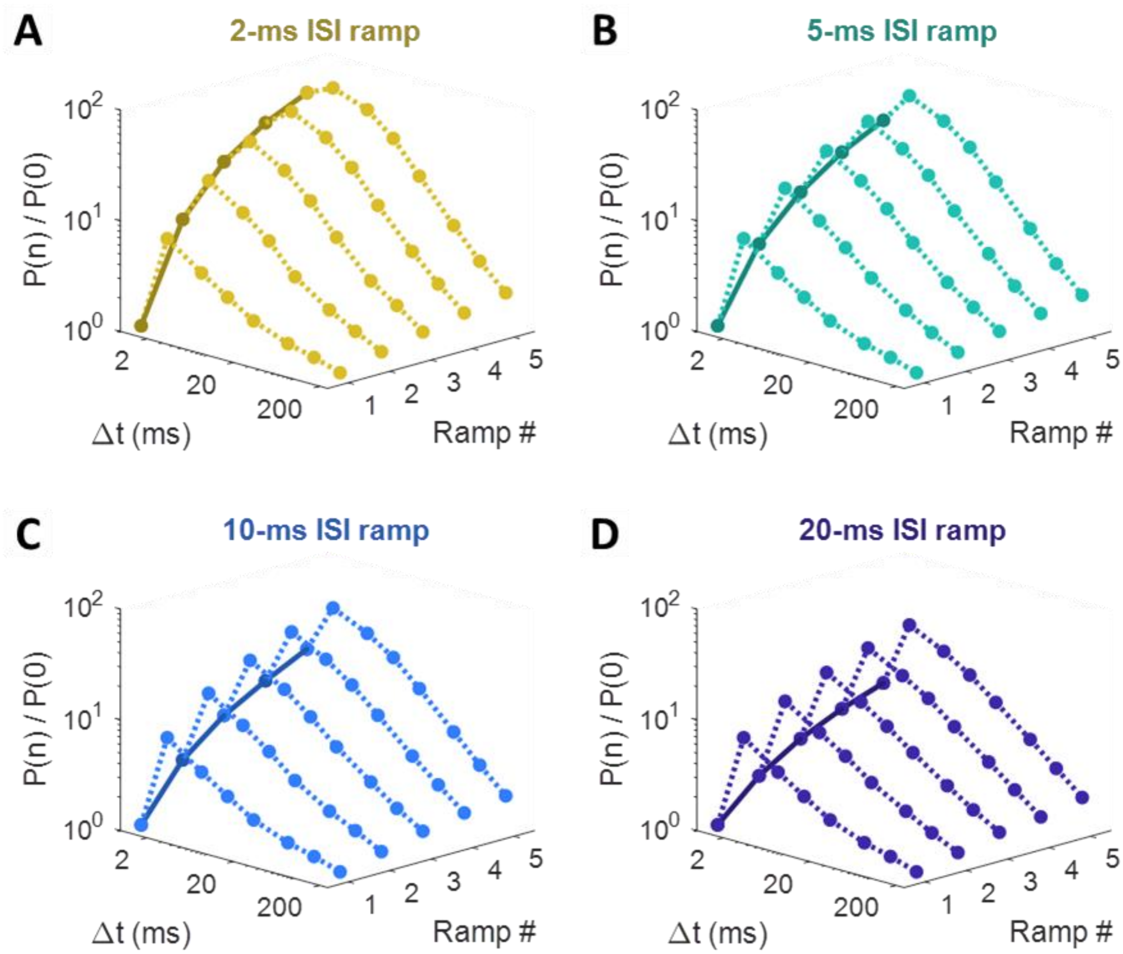


**S10 Fig. Empirical Facilitation in Release Probability is a Nonlinear Function of Spike History.**

Integrated release fidelity (P(n)) relative to baseline (P(0)) for the various stimulus cases explored. Ramp # indicates the number of spikes in the ramp preceding the probe spike, and Δt represents the ISI between the last ramp spike and the probe spike. Spike history noticeably affects the growth of facilitation, as seen for ramps with 2-ms ISIs (A), 5-ms ISIs (B), 10-ms ISIs (C), and 20-ms ISIs (D). Different colors distinguish facilitation functions with different spike histories. Dark lines follow relative release fidelity for spikes along spike ramps, and dotted lines follow relative release fidelity for probe spikes.

### Intuitive Exploration of Facilitation Function Behavior

The facilitation function introduced in this paper includes a nonlinear component that prevents the facilitation factor from exceeding some saturation limit. It does so by setting a limiting number of equal-sized steps to saturation, $N$, and then decreasing each step size according to

$$f(n) = f(n-1)\mathrm{e}^{-\Delta t/\tau} + 1 - \left(\frac{f(n-1)\mathrm{e}^{-\Delta t/\tau}}{N}\right)^N, \qquad F(n) = f(n)^\xi, \tag{40}$$

240 where $f(n)$ is the facilitation function after $n$ action potentials, $\Delta t$ is the time since the previous spike, $\tau$
241 is the facilitation decay time constant, $\xi$ is the facilitation factor nonlinearity, and $F(n)$ is the facilitation
242 factor to be multiplied by the release rate profile (see Eq (12)). Note that subscripts have been omitted
243 to focus on the facilitation of a single release component.

244 To gain a better intuition of this function, consider the limit as $N \to \infty$,

$$f(n) = f(n-1)\mathrm{e}^{-\Delta t/\tau} + 1, \qquad F(n) = f(n)^\xi. \tag{41}$$

245 As stimulus frequency becomes unphysiologically high ($\Delta t \to 0$), or as $\tau \to \infty$, the exponential decay
246 does not remove any facilitation between spikes and $f(n) = f(n-1) + 1$. In other words,

$$f(n|n \ll N) \approx n \quad \Rightarrow \quad F(n|n \ll N) \approx n^\xi \tag{42}$$

247 for sufficiently large $N$ and small $\Delta t/\tau$. Although $f(n)$ grows linearly for small $n$, the facilitation factor
248 eventually approaches its steady-state limit at

$$f(\infty) \approx N \quad \Rightarrow \quad F(\infty) \approx N^\xi \tag{43}$$

249 due to the third term of Eq (16) and (40). For cases with a smaller, constant frequency of stimulation,
250 the steady-state value for the facilitation component can be found by rearranging Eq (40) with
251 $f(n) = f(n-1) = f(\infty)$ and solving the polynomial

$$\left(N^{-N}\exp\left(-\frac{N\Delta t}{\tau}\right)\right) f(\infty)^N + \left(1 - \exp\left(-\frac{\Delta \mathrm{t}}{\tau}\right)\right) f(\infty) - 1 = 0. \tag{44}$$

252 For very large $N$, the first term approaches 0, yielding

$$f(\infty) = \left(1 - \exp\left(-\frac{\Delta t}{\tau}\right)\right)^{-1} \Rightarrow F(\infty) = \left(1 - \exp\left(-\frac{\Delta t}{\tau}\right)\right)^{-\xi}. \tag{45}$$

Thus, there is a finite, spike-frequency-dependent limit to facilitation even without the saturation parameter $N$. The function facilitates linearly for the first several spikes (for large enough $N$) and then plateaus to some maximum value. For large enough $N$ and $\xi = 1$, this set of functions acts as a simple convolution of an exponential with the spike times, so long as the $\tau$ of facilitation decay (Eq (16), (40)) exactly matches the $\tau$ of release rate decay (Eq (1), (3)). This kind of linearity, however, is not observed in the release profiles studied in this paper. S11 Fig A shows how different values for $N$ cause the otherwise linear step sizes to saturate at different levels. Importantly, $N \geq 1$ ensures stable growth. S11 Fig B shows how spike frequency also plays a role in determining the steady-state level of facilitation.

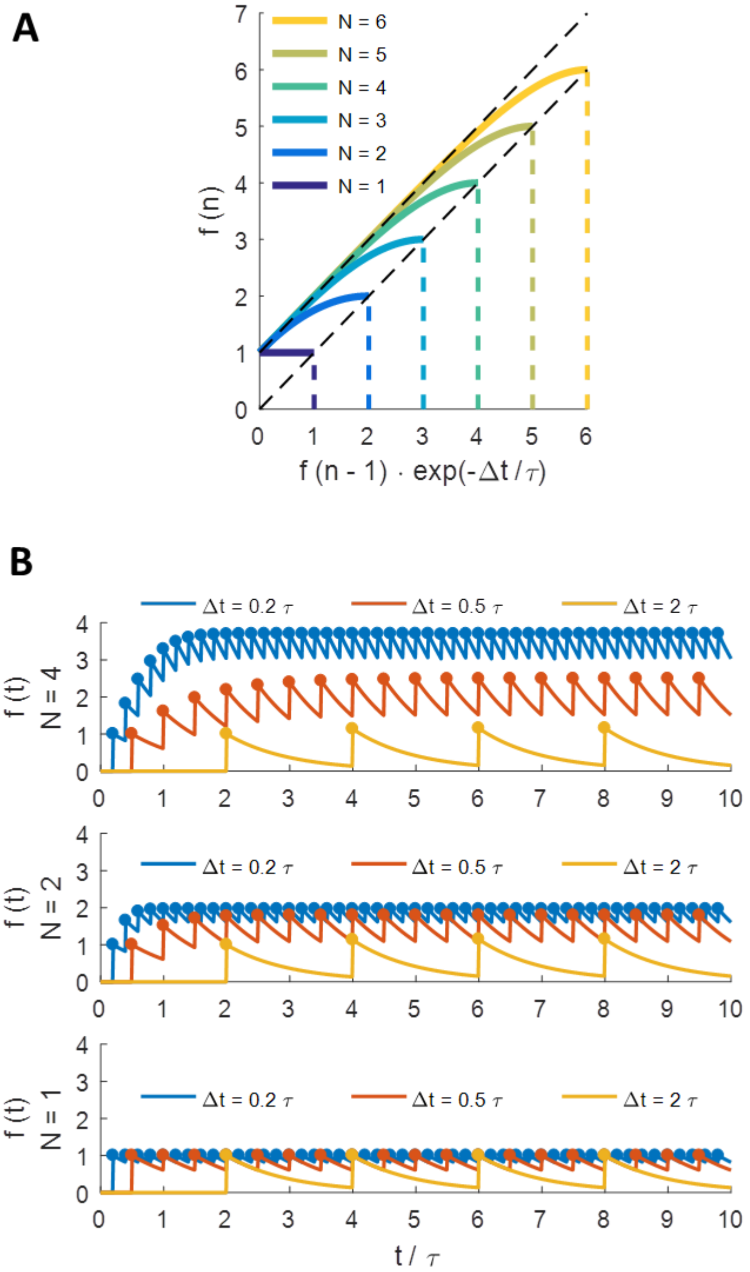


**S11 Fig. Saturation of Facilitation Parameters.**

A: Facilitation parameter f(·) increases almost linearly from one spike (f(n-1)) to the next (f(n)), until it approaches some limit N≥1. B: Curves represent the unseen change in f(·) between spikes. Dots represent actual values observed at spike times, values determined by the $Ca^{2+}$-triggered increment in release fidelity at each spike. Steady-state value for facilitation parameter limited by stimulus frequency and by value of N. No facilitation above baseline occurs for N=1.

Although we have considered facilitation always to be positive, this model provides the flexibility to allow negative facilitation. Whereas $F(\infty) > 1$ for $\xi > 0$, giving positive facilitation as normal, using $\xi < 0$ causes $F(\infty) < 1$, producing depression in the parameter. For $\xi = 0$, $F(\infty) = 1$, and no change can occur in the release-rate parameter. Such negative facilitation, although not observed in the magnitude of release rate for the Syt-1/7 mechanisms studied here, could apply in other circumstances to other parameters like time constants or rates that decrease with activity. For instance, short-term depression induced by $Ca^{2+}$-triggered inactivation of $Ca^{2+}$ channels [62, 88-90] could be represented as second or third component of the facilitation function that has a negative value for $\xi_{ci}$. However, this feature was not included in the MCell simulations, so it is beyond the scope of the current paper.

### Goodness of Fit of Facilitation Models

S12 Fig shows the fraction of the variance of the fitting error unexplained by the facilitation model (FVU error) for the final spikes of all 136 unique spike trains. Note how the highest error occurs with synchronous facilitation for interspike intervals (ISI) of 5 ms. More extensive exploration of facilitation space (i.e., longer spike trains with more diversity of spiking patterns) could elucidate an improved facilitation model that can achieve lower FVU error across all cases.

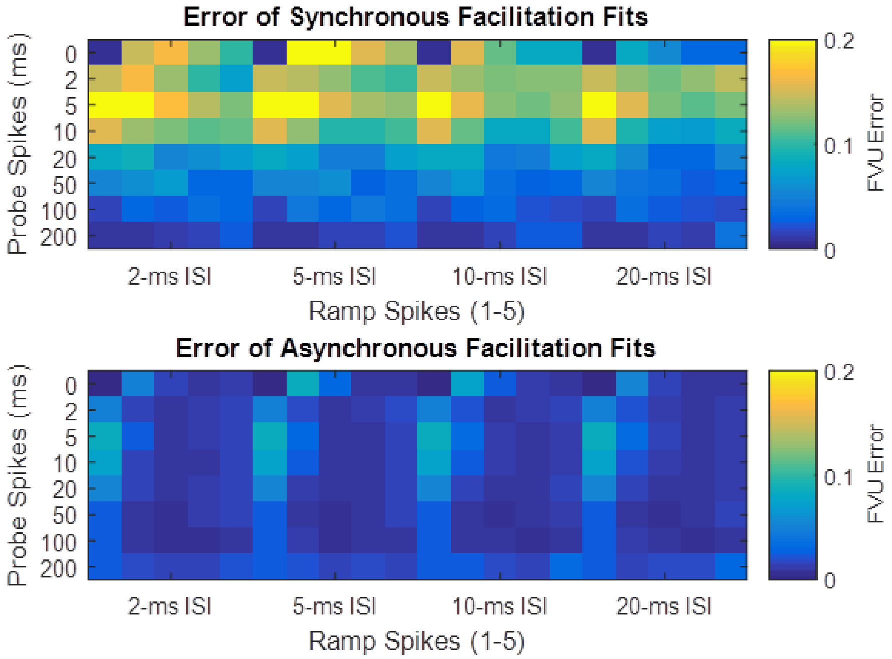


**S12 Fig. Release Rate Parameters and Facilitation Metaparameters Fitted to Empirical Histogram Profiles.**

Errors across all cases in linear and logarithmic space for the predictive model.
